# Linking plasmid-based beta-lactamases to their bacterial hosts using single-cell fusion PCR

**DOI:** 10.1101/2021.01.22.427834

**Authors:** Peter J. Diebold, Felicia N. New, Michael Hovan, Michael J. Satlin, Ilana L. Brito

**Affiliations:** Meinig School of Biomedical Engineering, Cornell University, Ithaca, NY; Robert Wood Johnson Medical School, New Brunswick, NJ; Weill Cornell Medicine, Cornell University, New York, NY

## Abstract

The horizonal transfer of plasmid-encoded genes allows bacteria to adapt to constantly shifting environmental pressures, bestowing functional advantages to their bacterial hosts such as antibiotic resistance, metal resistance, virulence factors, and polysaccharide utilization. However, common molecular methods such as short- and long-read sequencing of microbiomes cannot associate extrachromosomal plasmids with the genome of the host bacterium. Alternative methods to link plasmids to host bacteria are either laborious, expensive or prone to contamination. Here we present the One-step Isolation and Lysis PCR (OIL-PCR) method, which molecularly links target ARGs with the bacterial 16S rRNA gene via fusion PCR performed within an emulsion. After validating this method, we apply it to identify the bacterial hosts of three clinically relevant beta-lactamases in a neutropenic patient population who are particularly vulnerable multidrug-resistant infections. We detect novel associations of two low-abundance genera, *Romboutsia* and *Agathobacter*, with a multi-drug resistant plasmid harbored by *Klebsiella pneumoniae*. We put forth a robust, accessible, and high-throughput platform for sensitively surveying the bacterial hosts of mobile genes in complex microbial communities.

## Introduction

The emergence of multidrug-resistant (MDR) pathogens is a grave public health threat that occurs when pathogenic bacteria acquire antibiotic resistant genes (ARGs) through horizontal gene transfer (HGT) with bacteria in their proximal environment. The gut microbiome harbors a diverse repertoire of ARGs and these genes have been proposed to serve as a reservoir for HGT with MDR pathogens^1^. ARGs are often carried on mobilizable plasmids that impose technical challenges to surveying the set of bacteria affiliated with these genes. Standard molecular tools such as PCR and next-generation sequencing often fail to associate mobile ARGs with their bacterial hosts because they cannot capture the cellular context of extrachromosomal genes in the case of plasmids. Novel untargeted sequencing methods, such as bacterial Hi-C^2^ and methylation profiling^3^, provide broad reconstruction of plasmid-host relationships in metagenomes, as a trade-off for sensitivity. Alternatively, single-cell whole genome sequencing offers an ideal solution to this problem, but may be lower throughput, more expensive and require specialized equipment^4,5^. Targeted methods, such as bacterial cell culture under antibiotic selection, require that the ARG is expressed, functional, and selective in all hosts. Applying this broadly to capture the full diversity of the gut microbiome is complicated by the need for wide-ranging media and growth conditions^6,7^.

Single-cell qPCR is a targeted method to identify the hosts of specific genes, however each use specialized microfluidic devices, are limited in bacterial taxa they can capture, and most do not allow direct sequencing of the PCR products^8–10^. Alternatively, epicPCR^11^, uses fusion PCR and two emulsion steps to associate a taxonomic marker with a functional gene. Sequencing the fused PCR products provides accurate and sensitive associations between 16S sequence taxonomy and a given target gene. However, this method can be challenging to execute, difficult to scale up for multiple samples, and utilizes toxic and difficult-to-acquire reagents.

Here, we put forth One-step Isolation and Lysis PCR (OIL-PCR), a method that detects host-ARG associations from complex microbial communities through cellular emulsion and fusion PCR. Our streamlined method, based on the innovation of epicPCR, simplifies the procedure by combining the two emulsion steps of cell lysis and fusion PCR into a single emulsion PCR reaction that can be performed in a 96-well format using robotic automation. Furthermore, OIL-PCR can be multiplexed to target at least three genes in the same reaction, uses non-toxic commercially available reagents, and can be performed without relying on microfluidics or specialized equipment. Validation experiments on three environmental bacterial communities reveal that OIL-PCR is highly accurate and specific. We demonstrate the utility of this approach in examining the novel association of three extended spectrum beta-lactamase (ESBL) genes with two commensal organisms in the gut. Our results highlight the utility of this method in defining mobile ARG distribution within microbial communities as complex as the human gut microbiome.

## Results

### Development of a One-step Isolation and Lysis PCR method

OIL-PCR applies established fusion PCR methods to fuse any gene of interest to the 16S rRNA gene using three primers: two primers hybridize to the target gene, and a universal 16S reverse primer hybridizes to the V4 region. Amplification of the target gene appends a universal 16S forward primer sequence to the end of the target amplicon via a tailed reverse primer. The target gene amplicon then acts as a primer for amplification and hybridizes to the 16S rRNA gene as a forward primer, producing a fused gene product containing both the target gene and the 16S V4 sequence (Fig. 1a, Supplementary Fig. 1).

**Figure 1.**
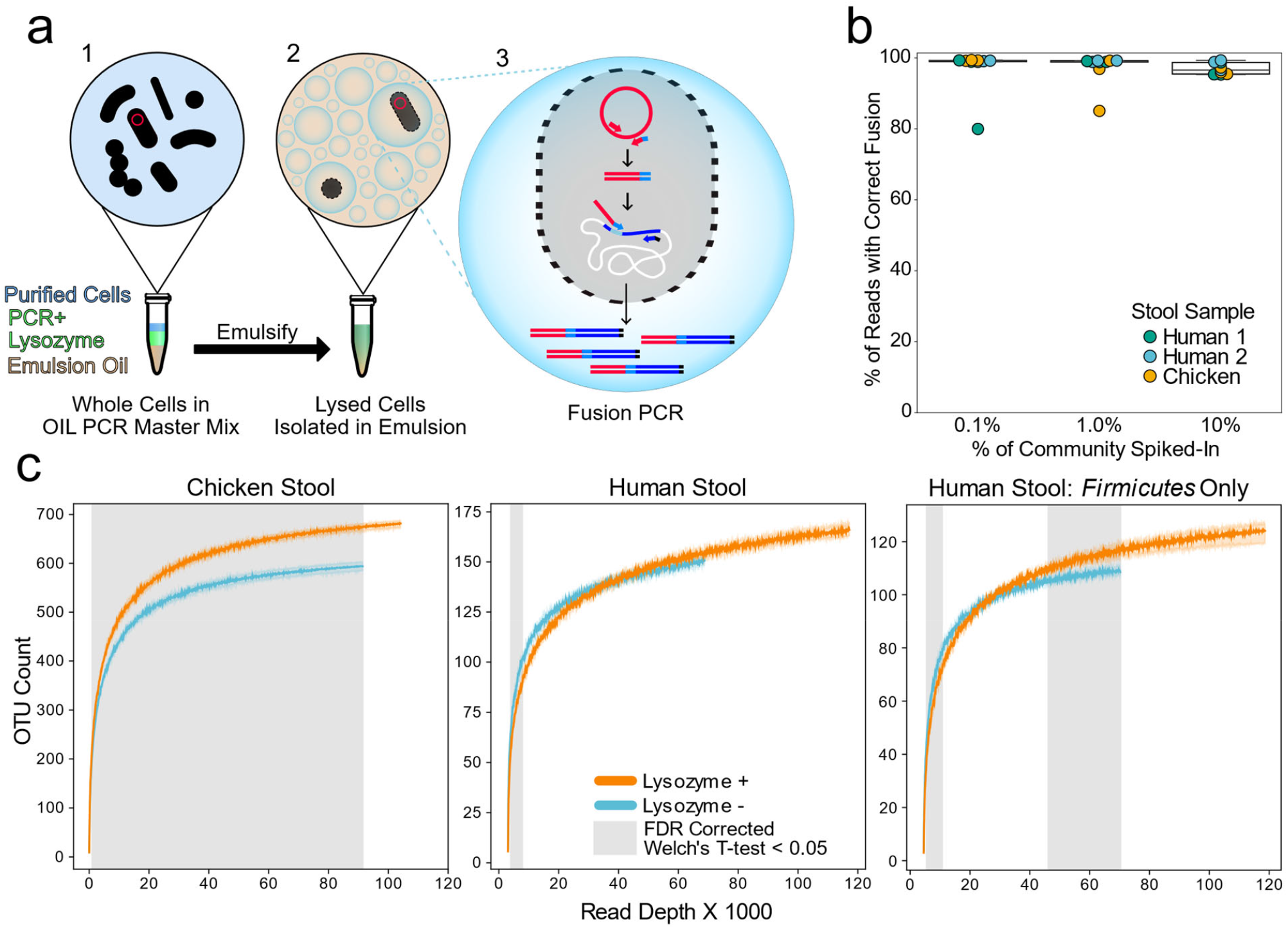
OIL-PCR can specifically link plasmid-encoded genes with their hosts. a) Depiction of the OIL-PCR method. (1) Nycodenz-purified cells are mixed with PCR master mix, lysozyme, and emulsion oil and shaken to create an emulsion. (2) Cells are lysed within the emulsion. (3) Fusion PCR is performed in droplets containing cells harboring the targeted gene. Fused amplicons between the gene of interest and the 16S rRNA gene are the product. b) A boxplot showing the percent of Illumina reads containing correct fusion products, namely the fusion of plasmid-borne *cmR* and the 16S rRNA gene of *E. coli* MG1655. OIL-PCR was performed on two individuals’ and one chicken’s gut microbiome sample, spiked with varying concentrations of *E. coli*. c) Rarefaction analysis of chicken (left) or human gut microbiome sample (middle) with (orange) and without (blue) lysozyme treatment. At right is the rarefaction analysis performed on Firmicutes only in the human stool sample. Grayed regions in the plot represent areas where the curves are significantly different (p<0.05) from one another, according to an FDR-corrected Welch’s t-test.

For fusion PCR to accurately link target genes with host marker genes, cells must be isolated to prevent the formation of non-specific fusion products. Oil emulsions and microwells have long been used to isolate eukaryotic cells, however, it is difficult to lyse bacteria in this format, especially gram-positive bacteria due to their thick cell walls. Existing single-cell isolation methods for bacteria either do not address this problem^9,10^, rely on specialized microfluidics^12^ or use time-consuming methods to encapsulate bacteria within hydrogel beads before performing multi-step chemical and enzymatic lysis procedures^11,13^. To address this problem, OIL-PCR combines bacterial isolation, lysis, and fusion PCR into a single streamlined reaction.

We developed a protocol that allows for the incorporation of Ready-Lyse (RL) Lysozyme into the fusion PCR master mix. Whole bacterial cells are added directly to the master mix while on ice to inhibit lytic activity during sample preparation. Vigorous shaking of the mixture then encapsulates the individual cells in an emulsion. Warming the emulsion to 37 °C activates the enzyme, lysing the cells. Next, a standard PCR thermocycler carries out the fusion PCR reactions in the single-cell emulsions. Fused PCR products are purified from the emulsion and amplified further with a nested primer to filter out off-target PCR products and add Illumina adapters. Lastly, custom indexing primers are used to index the fused products before Illumina sequencing. Our experiments confirmed the compatibility of the RL Lysozyme with the fusion PCR reaction, but required the addition of bovine serum albumin, a globular protein known to reduce protein aggregation^14^ (Supplementary Fig. 2a). We found that RL retained full activity in the standard NEB Phusion HF buffer (Supplementary Fig. 2c).

Next, we optimized the fusion PCR master mix to maintain a stable emulsion and amplify efficiently in picoliter droplets. PCR emulsions were prepared with fluorinated oil as used in modern emulsion-based methods, such as Drop-Seq^15^. We combined the fusion PCR master mix with bacterial cells and emulsion oil in either a 1.5 ml tube or a 0.5 ml deep-well plate before emulsifying the mixture using a tabletop bead homogenizer. Unlike microfluidic-enabled emulsions, our protocol leverages equipment commonly found in most molecular biology laboratories. We stabilized the emulsion by using detergent-free buffers and improved the efficiency of the PCR amplification within the emulsion by adding additional polymerase, BSA, dithiothreitol (DTT), and ammonium sulfate. We found that the addition of extra MgCl_2_ mitigated the inhibitory effects of extremely high concentrations of cell debris within droplets after lysis (Supplementary Fig. 1b).

### OIL-PCR accurately associates plasmid genes with the host in a binary community

In any emulsion-based method, it is essential to optimize the concentration of input cells to prevent the encapsulation of two or more cells in the same droplet. When using a monodisperse emulsion such as those generated using microfluidics, the ideal concentration of input cells is chosen using a Poisson distribution^8–10^. However, these calculations are not reliable in the case of a polydisperse emulsion, as employed here. We therefore developed a probe-based TaqMan qPCR assay to experimentally verify the optimal concentration of input cells that prevented non-specific gene fusions (Supplementary Fig. 3a). OIL-PCR was performed on a binary community consisting of *E. coli* carrying the chloramphenicol resistance gene *cmR* on a plasmid and WT *V. cholerae*. The two strains were mixed 1:1 and we performed OIL-PCR with a fusion primer set specific to *cmR* and universally targeting the 16S rRNA gene^11^ (Supplementary Table T2 and T3). A gradient of cell input concentrations was used and the final PCR products were recovered and purified. We then performed probe-based qPCR on the purified product using a nested primer for *cmR*, two blocking primers to inactivate any unfused amplicons, and two distinct fluorescent TaqMan probes (Thermo-Fisher 4316034) to specifically target the V4 region of either *E. coli* or *V. cholerae* (Supplementary Table T2). The fluorescent signal from each probe measured the relative ratio of specific to non-specific gene fusions present in the final amplicon pool (Supplementary Fig. 3a). When the input concentration of cells was at or lower than 400 cells/μl, or 40 k cells per reaction, non-specific gene fusion detection was reduced to undetectable levels (Supplementary Fig. 3b). As well as confirming that bacterial cells were isolated within the emulsion, we further confirmed that droplets did not coalesce by performing the TaqMan assay on OIL-PCR products from *E. coli* and *V. cholerae* cells combined after they were individually emulsified (Supplementary Fig. 3b). Our results confirmed that the emulsion is highly stable and coalescence was undetected.

### Application of OIL-PCR to environmental microbial communities allows robust and sensitive association of extrachromosomal elements with their host

Using OIL-PCR on environmental microbial communities requires clean bacterial cell preparations free of environmental contaminants which may inhibit PCR. To address this concern, cells were purified using Nycodenz density gradient centrifugation^16,17^, a simple method that can isolate clean bacterial fractions with minimal handling time to reduce contamination. Additionally, concerned that cell-free DNA can stick to the membranes and cell walls of bacteria^18^ thus introducing noisy associations in the data, we treated cells with heat-liable double strand-specific DNase (dsDNase). This enzyme only digests unprotected double stranded genomic DNA present in the samples without degrading single strand primers. By controlling the enzyme concentration, temperature, and speed at which cells were processed, we were able to digest extra-cellular DNA without impacting PCR efficiency of cellular contents. Using our Taqman assay, we demonstrated that including dsDNase treatment has the potential to increase the total cell input per reaction tenfold (Supplementary Fig. 3d).

To test the accuracy of our method on environmental samples, we spiked *Escherichia coli MG1655*^19^ containing plasmid pBAD33^20^ harboring *cmR* into two human and one chicken stool samples that lacked the gene according to PCR screening. Our results show that when *E. coli* was incorporated at 0.1%, or about 20 cells total, 97.8% of the reads (or 99.2%, excluding a single outlier) demonstrated the correct association when the test strain of *E. coli* was incorporated at 0.1%, or about 20 cells total, highlighting the sensitivity of OIL-PCR to detect the associations of genes in low abundant species across different sample types. The accuracy of OIL-PCR decreases slightly when the targeted sequence increases to 10% of the community composition, although associations were still 97% correct on average.

### Lysozyme improves capture of difficult-to-lyse gram positive bacteria

To achieve our goal of robust lysis and amplification to screen all bacteria within a complex community, we measured the effect lysozyme had on bacterial detection. We performed standard 16S sequencing on human and chicken stool communities using OIL-PCR, testing three variables: the effect of lysozyme, dsDNase, and heat inactivation of dsDNase on total bacterial recovery (Supplementary Fig. 4). All eight combinations of the three variables were tested in duplicate for two stool samples using robotic automation. For our analysis, we chose to focus on the total number of operational taxonomic units (OTUs) captured in our data rather than relative abundance metrics, as this better reflects our goal of detecting species, rather than recapitulating the starting community structure.

First, we assayed how each of the three variables (RL, dsDNase, and heat inactivation) affected OTU recovery. Based on rarefaction curves, we found dsDNase and Heat inactivation had no significant effect on OTU recovery in human and chicken stool, while RL lysozyme significantly increase OTU recovery in chicken stool based on Welch’s T-test with Benjamini-Hochberg FDR correction (Fig 1, Supplementary Fig. 4). RL was the only variable that significantly changed OTU recovery and therefore it was the only variable included in our final protocol.

Next, we looked to see whether any taxonomic groups were being enriched or depleted with lysozyme. Technical replicate OTU tables were combined to allow for deeper sampling depth. Results show that no phylum was significantly depleted, and importantly, at higher sequencing depths, *Firmicutes* were enriched in both the human and chicken samples. Additionally, *Bacteroides* species were enriched in the chicken sample. These results demonstrate the benefit of RL Lysozyme for capturing difficult to lyse gram-positive bacteria in the Firmicutes phylum, which account for much of the commensal diversity within the gut microbiome

We noticed that the total number of OTUs recovered from OIL-PCR was significantly lower than 16S sequencing of the input community at the same sampling depth (Supplementary Fig. 5). We hypothesized the reason for this dramatic reduction in OTUs was due to subsampling bias introduced through low cell input and variable amplification efficiency in OIL-PCR. To test our hypothesis, we combined OTU tables from two, four and eight technical replicates and found a consistent up-shift for each rarefaction curve as we combined more tables. This up-shift was not observed when combining the input Nycodenz sequencing, indicating that the reduced OTU counts were due in part to sub-sampling bias and not an inherent failure to capture bacterial taxa (Supplemental Fig. 5). We therefore recommend OIL-PCR to be performed in replicates such that a sufficient number of cells are being sampled.

### Increased throughput through automation and multiplexing

To further improve the efficiency and throughput of OIL-PCR, we sought to transition the method from 1.5 ml centrifuge tubes to a 96-well plate format using the Eppendorf epMotion liquid handling robot. The liquid handling robot can perform certain parts of the PCR preparation as well as DNA recovery and purification. The automated workflow allowed us to process up to 48 samples simultaneously with fewer manual steps overall.

We next tested whether OIL-PCR could simultaneously target multiple genes though multiplexing. We repeated the previously described TaqMan assay using a strain of *V. cholerae* containing the ampicillin resistance gene *ampR* and *E. coli* with *cmR;* both on a plasmid (Supplementary Fig. 3c). Our results demonstrate that OIL-PCR can be multiplexed while still accurately maintaining the correct associations of target genes with their host bacteria.

### Bacterial hosts are identified for several clinically important ß-lactamase genes

We analyzed metagenomic sequencing of stool samples that were collected from a cohort of patients who were neutropenic because of chemotherapy administered for a hematopoietic cell transplant. Two patients, B335 and B314, were chosen for OIL-PCR based on the presence of three class-A beta-lactamase genes, *bla*_*TEM*_, *bla*_*SHV*_, and *bla*_*CTX-M*_ in the metagenomes. We tested a three-sample time course from patient B335: before antibiotic treatment, after four days of trimethoprim-sulfamethoxazole and one day of levofloxin, and lastly after an additional two days of levofloxin (Fig. 2a). Patient B335 carried all three genes across three time points with *bla*_*TEM*_ and *bla*_*CTX-M*_ on an 80 kb *Klebsiella* plasmid and *bla*_*SHV*_ on a contig that was annotated as *Klebsiella* (Fig. 2b). We tested one sample from patient B314 from before antibiotic treatment which carried multiple *bla*_*SHV*_ genes(Supplementary Fig. 6). We hypothesized that OIL-PCR could be used to sensitively and accurately detect additional hosts of these genes.

**Figure 2.**
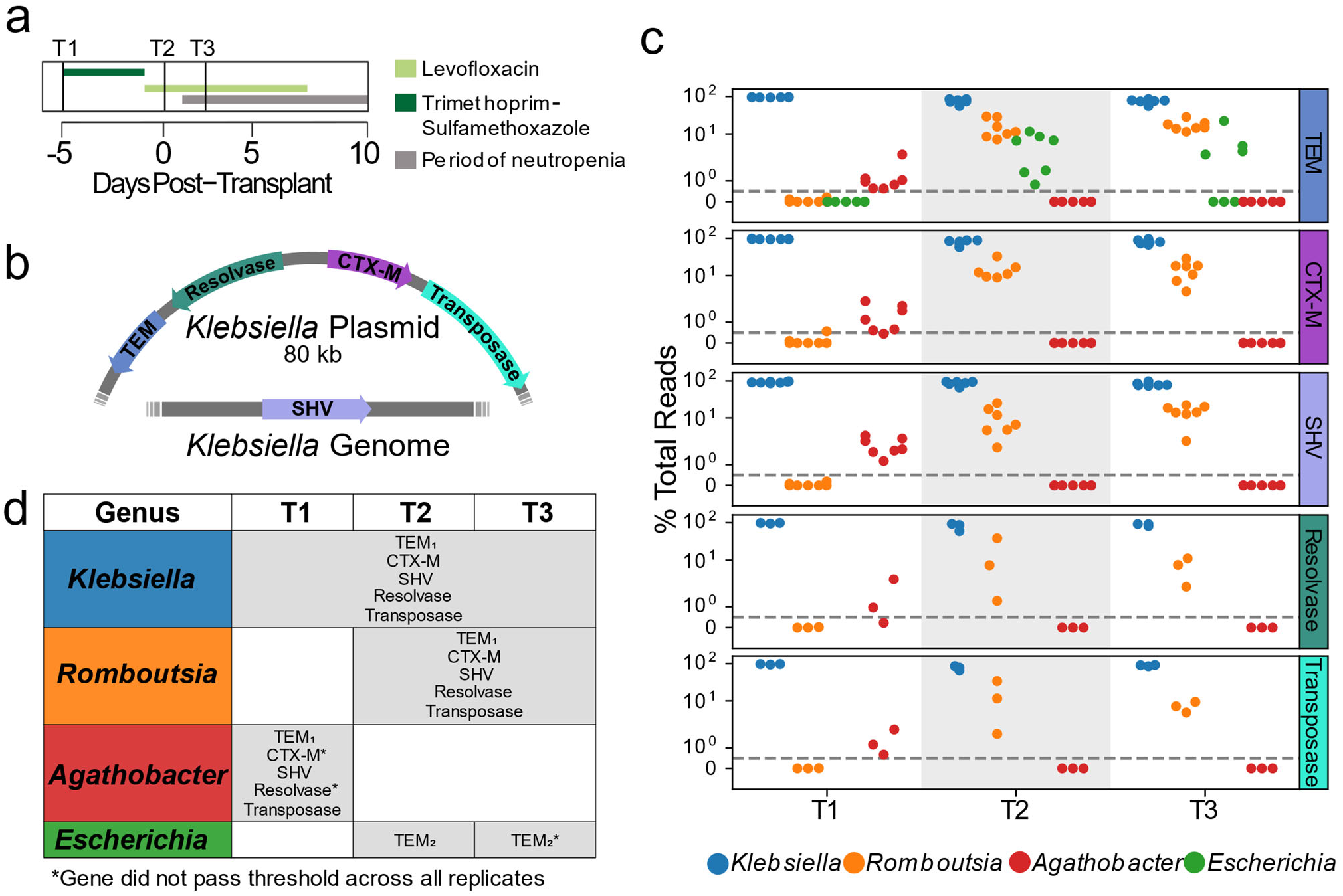
Extended spectrum beta-lactamase genes are associated with both pathogenic and commensal species. a) Summary of treatment and sample time points for patient B335 b) Depiction of an 80kb plasmid carried by *K. pneumoniae* harboring the *bla*_*CTX-M*_, *bla*_*TEM*_, Tn3 transposase and resolvase genes. The *bla*_*SHV*_ gene is presumed to be carried within the *K. pneumoniae* genome. Placement of these genes was inferred from metagenomic assemblies of patient B335’s gut microbiome sample. c) OIL-PCR results for each of the genes depicted in (a) patient B335 at 3 time points. For all gene-taxa associations, the percent of total OIL-PCR reads for that gene-time point is plotted. All species passing our detection threshold of 0.5% (dotted line) at any of the three time points is included in this plot. d) A table summarizing the results in (b). All gene-taxa associations for each time point passing our detection thresholds are listed. Two SNP variants of TEM were detected and denoted with subscript numbering. Gene-taxa associations which did not consistently pass our detection threshold across all replicates are noted (*).

We designed three degenerate fusion primer sets to broadly target most variants of *bla*_*TEM*_, *bla*_*SHV*_, and *bla*_*CTX-M*_ (Supplementary Table T2, T3), and performed multiplexed OIL-PCR with robotic automation. Samples were processed in quadruplicates along with negative and positive controls. We set a threshold for defining positive gene-taxa associations, as having 0.5% of total reads across the four replicates.

Our OIL-PCR results largely confirm findings in the metagenomic assemblies from Kent et al.^2^. In B314, we found *bla*_*SHV*_ associated with *Klebsiella* as suggested by metagenomic assemblies. However, we also detected two other class-A beta-lactamase genes, *bla*_*LEN*_ and *bla*_*OXY*_, which were present in the metagenomes but we did not expect to amplify with our primers. *bla*_*LEN*_ amplified with the primers designed for *bla*_*SHV*_ and *bla*_*OXY*_ amplified with primers for *bla*_*CTX-M*_. Curiously, *bla*_*OXY*_ is an exceptionally poor match for our *bla*_*CTX-M*_ primers, having a mismatch one base away from the 5’ end of the fusion primer. We hypothesize that the low annealing temperature and modified buffer used in the emulsion PCR is highly permissive to priming mismatches. We see permissive annealing as an advantage for the method because it allows for amplification of unknown variants of target genes while amplification due to off-target priming is filtered out during the nested PCR step (Supplementary Fig. 1), leaving only the true amplicons in the final sequencing. This permissive annealing behavior of OIL-PCR can be leveraged in the future to design broad-range primers for diverse gene groups such as metallo-beta-lactamases^24^.

Results from patient B335’s time course also matched the metagenomic sequencing from Kent et al., associating *bla*_*TEM*_, *bla*_*SHV*_, and *bla*_*CTX-M*_ with *Klebsiella* in all three time points (Fig. 2c,d). We also found that all three genes strongly associated with the commensal genus *Romboutsia* in time points T2 and T3 and to a lesser extent with *Agathobacter* in time point T1 (Fig. 2c,d). A strain of *Escherichia* with a distinct variant of *bla*_*TEM*_ was detected at time point T2, but did not pass the detection threshold across all replicates in timepoint T3. We repeated OIL-PCR on all three samples from B335, this time in triplicate without multiplexing to further confirm these results. The singleplex experiment perfectly mirrored the multiplex results, excluding one replicate of T2/CTX-M which failed to sequence, indicating that these genes are linked with organisms other than *Klebsiella*. As further confirmation of this result, we targeted two Tn3-like transposon genes situated in close proximity to *bla*_*TEM*_ and *bla*_*CTX-M*_ on the 80 kb *Klebsiella* plasmid. We hypothesized that these genes should also be associated with the same genera as the ARGs. Remarkably we observed the identical pattern with *Klebsiella, Romboutsia*, and *Agathobacter* as with the three beta-lactamases, but not *Escherichia*, which carried a distinct variant of *bla*_*TEM*_ (Fig 2c,d).

### OIL-PCR provides further evidence of the association of beta-lactamases with the commensal *Romboutsia*

We next investigated whether OIL-PCR could be used to further confirm the association between *Romboutsia* and the three beta-lactamases. We focused specifically on *Romboutsia* because of the strong signal in the OIL-PCR results compared to *Agathobacter*. For this experiment, instead of fusing the ARG sequence to the 16S rRNA gene using universal primers, we used primers designed to specifically detect the *Romboutsia* 16S rRNA^25^ and fused the 16S gene specifically to *bla*_*TEM*_ (Fig. 3a, Supplementary Table T2, T3). In this instance, no amplification is possible unless *Romboutsia* is encased in the same droplet with the *bla*_*TEM*_ gene and would negate the possibility of false-positive associations due to chimera formation. Results show amplification and sequencing was only produced from time points T2 and T3 with no signal detected at time T1, confirming the presence of *bla*_*TEM*_ within *Romboutsia* at time points T2 and T3 but not T1 (Fig. 3b).

**Figure 3.**
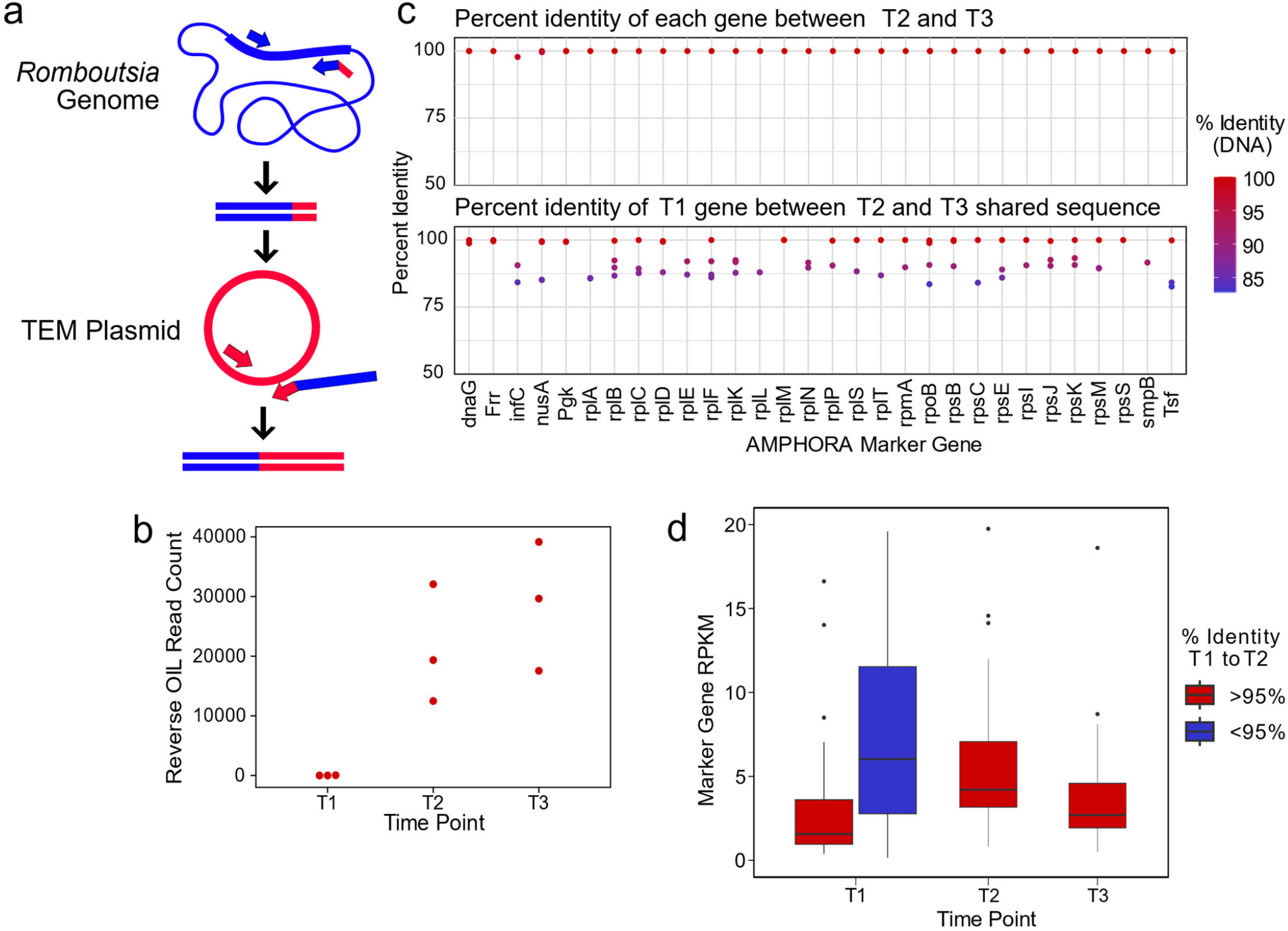
*R. timonensis* strains associated with the three beta-lactamase genes appear over the patient’s time course. a) Depiction of the reverse OIL-PCR reaction in which *Romboutsia*-specific 16S rRNA sequences (blue) are fused with the *bla*_*TEM*_ sequences (red). b) OIL-PCR read counts of the reaction shown in (a) are plotted. c) The percent sequence identity of assembled *R. timonensis* marker genes between genes identified in timepoints 2 and 3 (top) and between timepoints 1 and between sequences shared at timepoints 2 and 3 (bottom). d) RPKM-normalized abundance-values for the assembled marker genes for each strain assembled in time point 1 and the major strain present in timepoints 2 and 3.

We next explored the metagenomic data for clues as to whether the *Romboutsia* strain was present at timepoint T1, but below the detection threshold, or whether the strain linked with the genes was acquired sometime between time T1 and T2. Based on the 16S data from OIL-PCR and metagenomic sequencing, we identified the *Romboutsia* species as *R. timonensis*. Genus-level abundance data showed *R. timonensis* to be present in all three timepoints in patient B335. Due to the overall low abundance of this organism, we were unable to assemble a *Romboutsia* genome from these samples. Instead, we aligned patient B335’s three samples to the *R. timonensis* (PRJEB14233) genome from NCBI, assembled the aligned reads and examined similarities between the *R. timonensis* taxonomic markers over the three timepoints (Figure 3c). We found that B335 was colonized by at least two independent strains of *R. timonensis* during the first timepoint, but that only one *R. timonensis* strain persisted during timepoint T2 and T3. One of the *R. timonensis* strains from T1 was identical to the strain from T2 and T3 across 15/30 AMPHORA marker genes, and >99% identical in 24/30 genes (Fig. 3c), suggesting that the strain of *R. timonensis* from time T2 and T3 was also present at time point T1. We found no significant difference in the normalized abundance of *Romboutsia* between time point T1 and time points T2 and T3 (Fig. 3d), albeit our data suggests that the persistent strain is the minor variant at time point 1. Despite the sensitivity of OIL-PCR, which can detect cells at least 0.1% abundant (Fig. 1b), we cannot rule out the possibility that the stochasticity of sampling in OIL-PCR and the low abundance of this particular strain of *R. timonensis* precluded our ability to observe this association at the beginning of the time course.

## Discussion

Here we show the ease with which OIL-PCR can identify novel carriers of known resistance markers on extrachromosomal elements within complex bacterial communities. We applied it to a neutropenic patient’s microbiome and showed the correct association of three beta-lactamases with *K. pneumoniae*, and also discovered novel associations between these beta-lactamases and two gut commensals, *R. timonensis* and *Agathobacter* spp. Two of the genes, *bla*_CTX-M_ and *bla*_TEM_, were both found on a large *Klebsiella* plasmid within the metagenome, suggesting the possible transfer of these genes to *R. timonensis* during the time course. Analysis of the plasmid sequence showed that it contains an origin of transfer, but does not have the genes necessary to transfer itself, meaning it would require a second “helper plasmid” to mobilize. Additionally, *bla*_SHV_ was only found on a contig belonging to the *Klebsiella* genome without any known mobilizable transposons or integrative conjugative elements nearby, severely limiting its transfer potential. An alternative explanation for our results is that *Romboutsia* and *Klebsiella* became physically associated within the gut, and thus consistently emulsified together. Both of these explanations highlight the dynamic nature of the gut microbiome, either through horizontal gene transfer, or novel physical associations between pathogens and commensals, with close physical association being a known activator for conjugal transfer of genes^26^. In future applications of OIL-PCR, primers targeting non-transferrable genes could be used to distinguish between transfer and aggregation when identical genes are associated with different taxa.

Our results illustrate the application of a streamlined, simplified fusion PCR approach to obtain robust and sensitive associations of extrachromosomal DNA with bacterial hosts. It is a practical and transportable protocol with no requirements for specialized equipment nor specialized expertise. We identify improvements in performing single cell analysis on stool, namely the incorporation of a Nycodenz purification step and the incorporation of lysozyme plus heat-induced lysis. Additionally, we increased throughput at least three-fold through primer multiplexing and developed an automated protocol to process at least 48 samples concurrently, allowing a total of 144 gene-sample association tests per batch.

Additional improvements to OIL-PCR could be explored to further increase throughput and sensitivity. Although we tested multiplexing three genes per reaction, this number could likely be increased as we have found no sign of false positives due to multiplexing as demonstrated by associating a novel *bla*_TEM_ variant with only *Escherichia* in time point T2 of patient B335 (Fig 2b, c). Further, we show that the OIL-PCR master mix facilitates permissive annealing of primers, allowing a mismatch one base from the primer’s 3’ end as demonstrated when *bla*_OXY_ was detected in sample B324-2 with *bla*_CTX-M_ primers. These results could allow for the development of highly degenerate primers to target a broad range of gene variants. Non-specific priming during OIL-PCR is not of concern because the nested PCR reaction specifically filters out undesired fusion products. Lastly, the method described currently allows 40,000 cells total per reaction. However, our TaqMan based qPCR assays suggest that the input concentration could be increased 10-fold by pre-treating cells with dsDNase (Supplementary Fig 4d). Combined with our result showing that OIL-PCR is more accurate when detecting low abundant taxa (Fig 1b), we feel confident that cell input can be increased to improve sensitivity without sacrificing accuracy.

OIL-PCR is a highly versatile platform that could be applied across fields to address a multitude of questions. While we were interested in plasmid-born ARGs in the gut, the method could be used to target any gene of interest that is difficult to associate with a host using metagenomics. As mobile genetic elements are notoriously difficult to assemble due to their promiscuity which complicates de Bruijn graph assembly^27^, this method could be applied to find the hosts of integrated and non-integrated mobile elements. Similarly, as metavirome sequencing has revealed a massive number of viral genomes with unknown hosts^28^, OIL-PCR may be particularly useful in addressing this gap. Additionally, viral and plasmid host-range is an important determinant for understanding and modeling bacterial ecology of predation and HGT^29^. Further, targeting functional metabolic genes detected in metagenomes, but present at low abundance in bacterial communities, could identify novel bacteria involved in nutrient cycling which has remained a persistent challenge in the field of bacterial ecology^30^. Finally, when combined with microfluidics, direct lysis of bacteria in an emulsion, as shown here, could be used to develop or simplify single-cell genome sequencing or single-cell RNA-seq for bacteria.

## Supplemental Figures

**Supplemental Figure 1.**
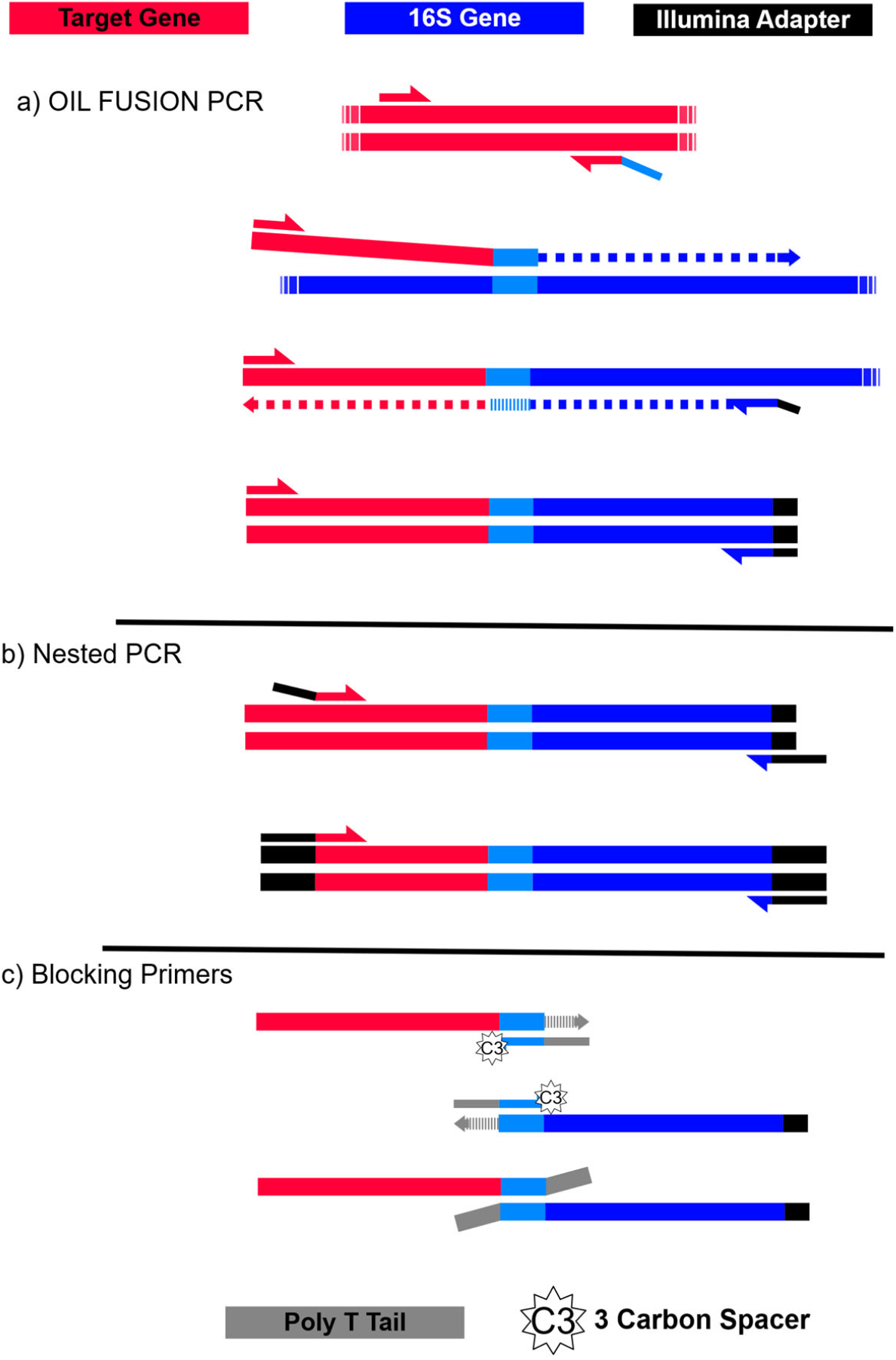
Depiction of the fusion PCR reaction. a) PCR is initialized with primers to a target gene (red). The reverse primer contains a 5’ overhang complimentary to the universal V4 16S primer 519F (light blue). The product of this first amplification step can act as a forward primer for the 16S rRNA gene (blue). After extension of the full fusion product, the forward target primer can pair with the universal 16S reverse primer 786R (with a portion of an Illumina TruSeq adapter sequence) to amplify the fully fused PCR product. b) Nested PCR is performed on the fused PCR products from (a) in order to filter out non-specific priming from the fusion PCR. The forward primer anneals within the target gene and has a TruSeq adapter at the 5’ end. The reverse primer also has the Illumina adapter sequence at it’s 5’; end and anneals to the non-degenerate portion of 786R and the partial Illumina adapter sequence appended in (a). c) Two blocking primers, both complementary to the 519F priming region, are included in the nested PCR to prevent unfused PCR products from annealing during the nested reaction. Blocking primers have a 3-carbon spacer on the 3’ end to prevent extension and a poly-T tail that appends 10 As to the 3’ end of any unfused products, thus inactivating them from annealing or extension.

**Supplemental Figure 2.**
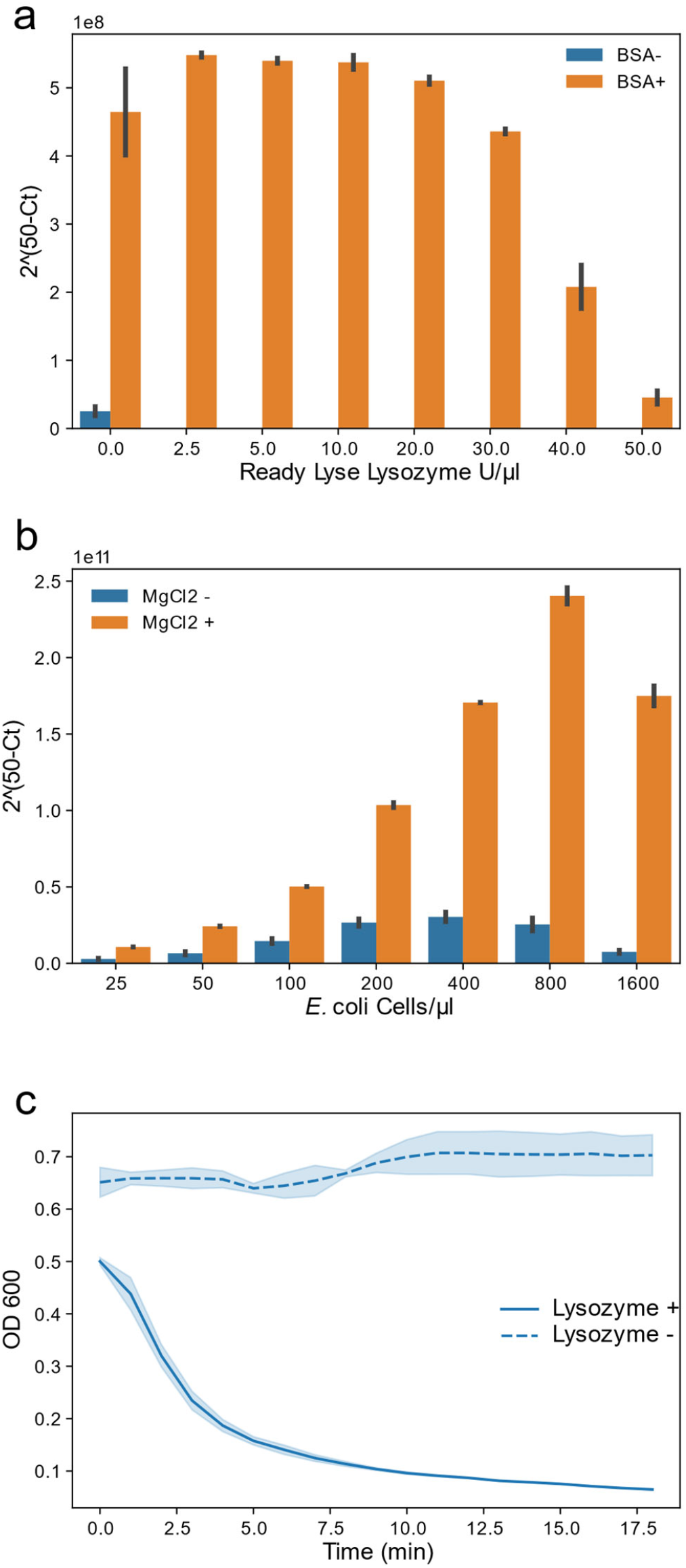
BSA and excess MgCl_2_ improve the efficiency of OIL-PCR and Ready Lyse Lysozyme remains active in OIL-PCR master mix. a) Sybr-based qPCR was performed on the *cmR* gene carried on pBAD33 with varying concentrations of lysozyme in the presence (orange) or absence (blue) of BSA. Higher 2^(50-Ct)^ values represent greater amplification. b) Sybr-based qPCR was performed on the *cmR* gene carried on the pBAD33 plasmid in *E. coli* MG1655 cells at increasing cell concentrations with (orange) and without (blue) additional MgCl_2_. Higher 2^(50-Ct)^ values represent greater amplification. c) Lysozyme activity against *B. subtilis* suspended in the OIL-PCR optimized reaction mix with (solid line) and without (dashed line) lysozyme.

**Supplemental Figure 3.**
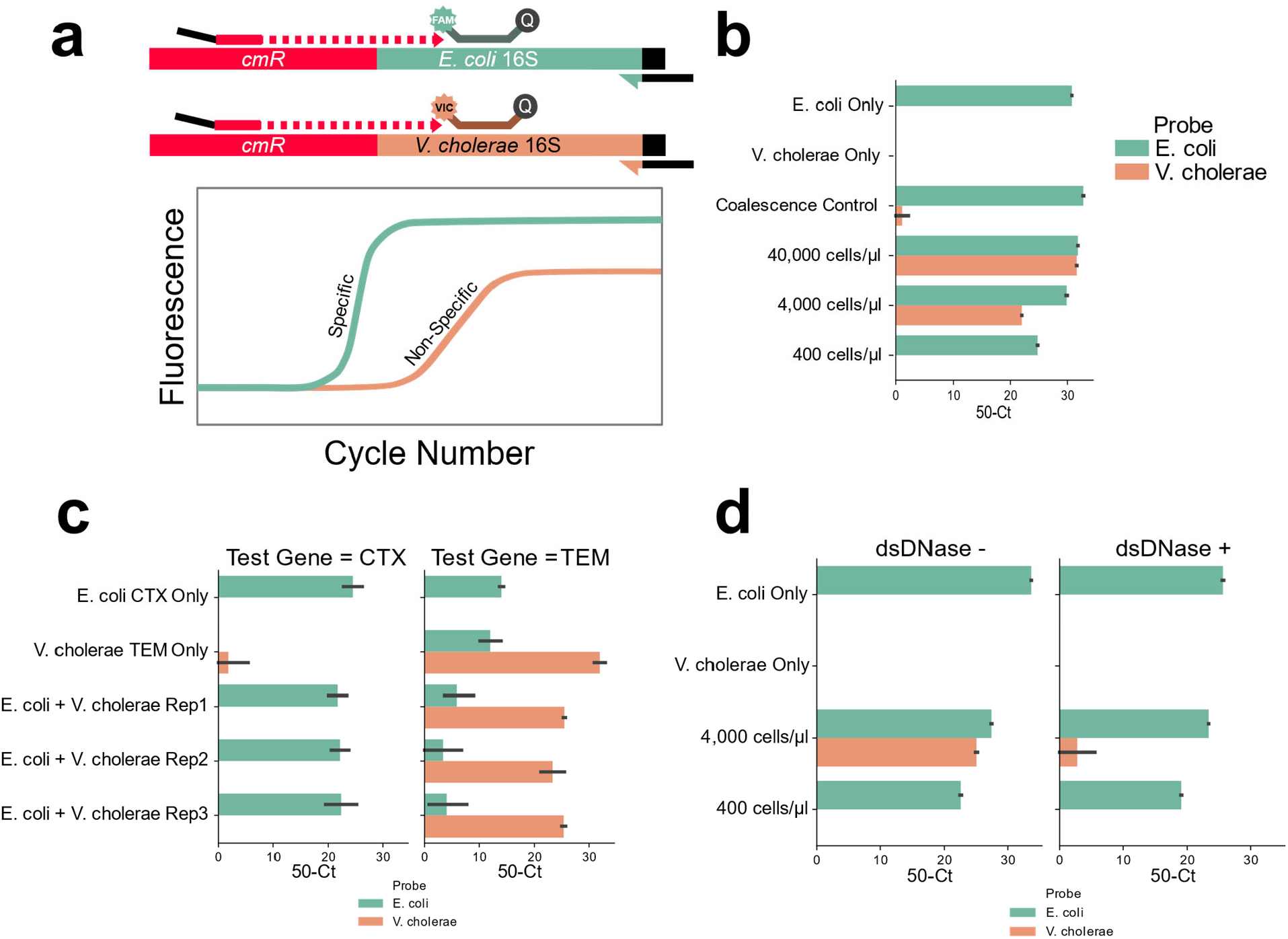
Cell concentration of 400cells/μl, DNase treatment, and multiplexing PCR reactions result in accurate OIL-PCR results. a) Diagram of the Taqman assay used to monitor OIL-PCR results. Briefly, Taqman probes were designed to be complementary for the 16S rRNA genes in either *E. coli* or *V. cholerae*, each with its own fluorophore. OIL-PCR was performed on *E. coli* carrying the *cmR* gene on the pBAD33 plasmid but not present in *V. cholerae*. Fusion PCR products were recovered and nested probe-based qPCR was performed. Upon amplification of the gene, the probe is cleaved by Taq polymerase releasing the fluorophore from the quencher. Specific amplification of the designated region is measured by fluorescence of the expected fusion product vs the non-specific product. b) OIL-PCR with primers targeting a plasmid-borne *cmR* gene was performed with a 1:1 mix of *cmR* positive *E. coli* and *cmR* negative *V. cholerae* cell suspensions with. A gradient of cell concentrations was tested (400-40,000 cells/μl), in addition to *E. coli* and *V. cholerae* suspensions alone as positive and negative controls. Control emulsions were mixed 1:1 after emulsification to test for droplet coalescence. c) Multiplexed OIL-PCR was performed with primer sets targeting a genomic *bla*_*CTX-M*_ gene in *E. coli* and a plasmid-borne *bla*_*TEM*_ gene in *V. cholerae*. Experiments were performed in triplicate and on each of the organisms separately. Results are shown for the *bla*_*CTX-M*_ (left) and *bla*_*TEM*_ (right). d) OIL-PCR with primers targeting a plasmid-borne *cmR* was performed after pretreating cells with (right) and without (left) dsDNase at two different 1:1 *E. coli* to *V. cholerae* cell suspension concentrations as well as on the individual bacterial strains for controls.

**Supplemental Figure 4.**
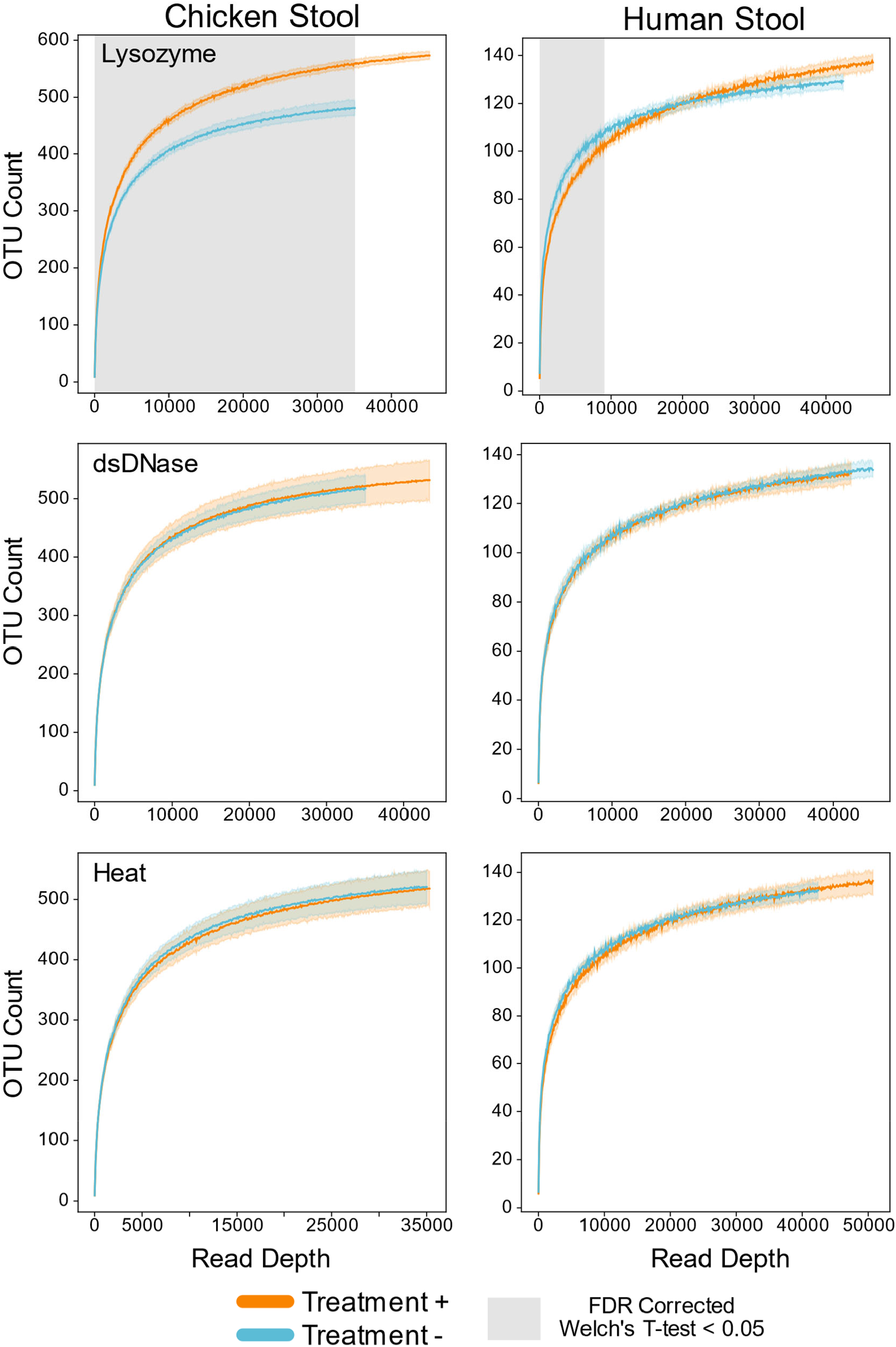
Lysozyme alone improved recovery of species. Rarefaction analysis of chicken (left) or human (right) gut microbiome samples with (orange) and without (blue) lysozyme (top), dsDNA treatment (middle) and heat (bottom). Grayed regions in the plot represent areas where the curves are significantly different from one another (p<0.05), according to an FDR-corrected Welch’s t-test.

**Supplemental Figure 5.**
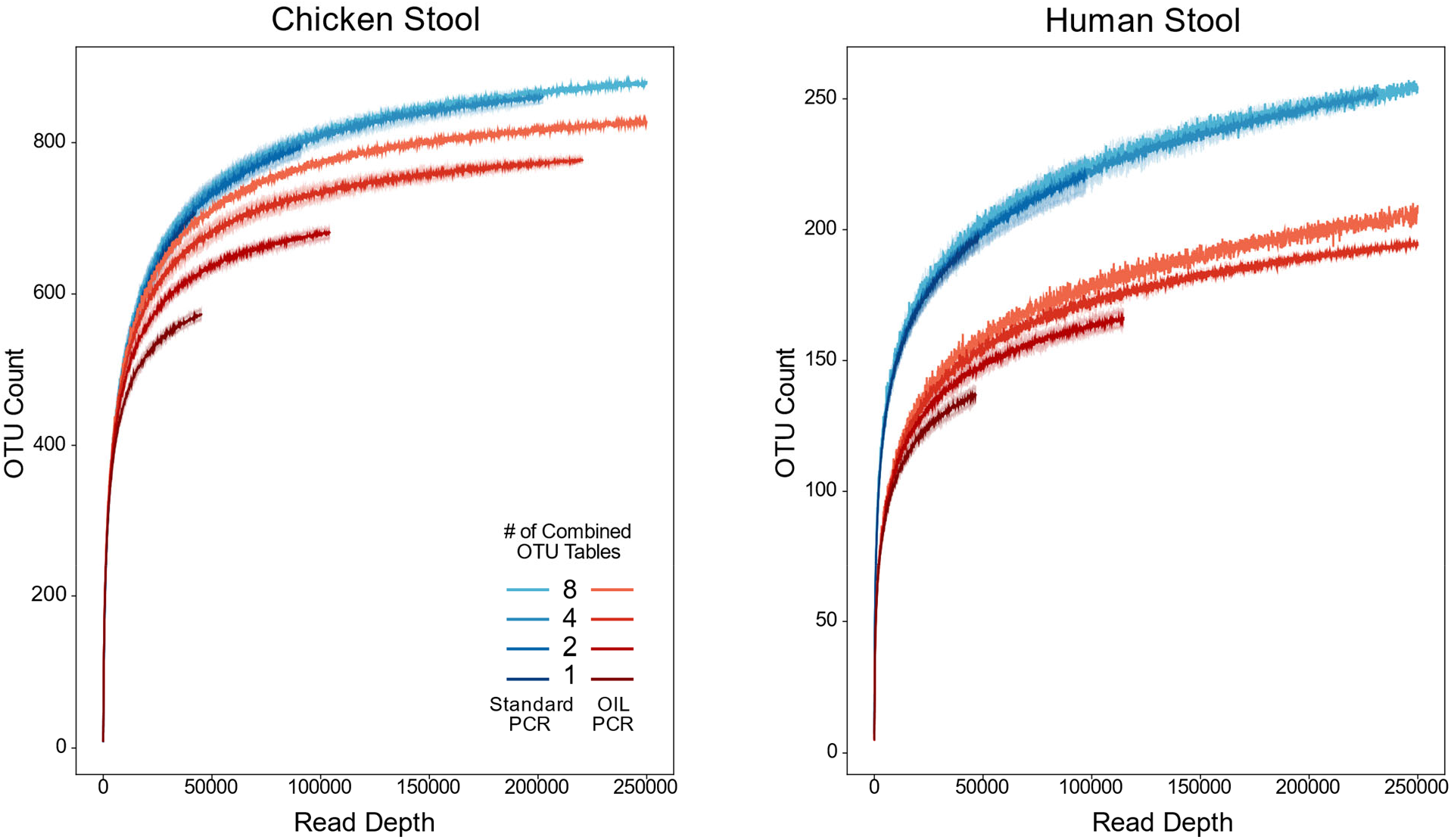
Combining replicates for increased depth improved recovery of species and reduced stochastic sampling bias. Rarefaction analysis of chicken (left) or human (right) gut microbiome samples performed with OIL-PCR (red) or standard DNA extraction and library prep (blue). OTU tables were repeatedly subsampled in groups of 2, 4, or 8 replicates from 8 total replicate 16S libraries.

## Methods

### Optimizing OIL-PCR Reaction Buffer for Phusion and RL Lysozyme Compatibility

#### SYBR-Based qPCR Assay for Phusion Polymerase Activity with Lysozyme

SYBR-based qPCR reactions were set up in duplicate as follows: 25 μl reactions with 20 U/ml Phusion Hot Start Flex DNA polymerase (NEB M0535L), 1X HF Buffer, 200 μM dNTP mix (NEB N0447L), 400 nM of 519F and 786R, 1X SYBR Green (Thermo Fisher S7563), 1X ROX reference dye (Thermo Fisher 12223012), 0.5 mg/ml of BSA (NEB B9000S) when included, 0.01% Triton-X 100, and 1μl of template DNA. Reactions were prepared with 0, 2.5, 5, 10, 20, 30, 40 and 50 U/μl of Ready-Lyse Lysozyme (Lucigen R1810M). qPCR was performed on the Thermo Fisher Quant Studio 3 Real-Time PCR machine with the following parameters: 98 °C for 1 minutes, then 50 cycles of 98 °C for 5 seconds, 54 °C for 30 seconds, and 72 °C for 30 seconds. Proper amplification was confirmed using melt curves: 98 °C for 5 seconds, cool to 60 °C at 1.6 °C/s, and then heat to 95 °C at 0.15 °C /s. Ct values and melt curves were generated with the Quant Studio software V1.4 using the default software settings.

#### Lysozyme Activity Assay in OIL-PCR Master Mix

Lysozyme testing was performed in a lysozyme test buffer made from the OIL-PCR master mix with dNTPs, primers, and Phusion polymerase replaced with water and 100% glycerol (48ul/ml). Log phase cultures of *B*. subtilis were standardized to an OD_600_ of 2 and suspended in 1x Lysozyme test buffer. Separately, lysozyme was suspended in 1x Lysozyme test buffer at 2x concentration. 100μl of the lysozyme mix was aliquoted into a 96-well, clear, flat-bottomed microtiter plate before adding 100μl of suspended culture. Lysis was monitored using a Spectramax M3 plate reader (Molecular Devices), heated to 37 °C and OD_600_ measured every minute for an hour.

#### Optimizing OIL-PCR Reaction Efficiency in an Emulsion

50μl PCR reactions were prepared in 1.5 ml tubes as describe in the Tube-Based OIL-PCR method below with varying concentrations of Phusion polymerase, bovine serum albumin (BSA), Ready Lyse lysozyme, dithiothreitol (DTT), MgCl_2_, dNTPs, and ammonium sulfate. *E*. coli genomic DNA was used as template and amplified with universal 16S primers 519F and 796R. Reactions were emulsified 25 Hz for 30 seconds before aliquoting to PCR tubes and thermocycling. Final PCR products were separated from the emulsion as describe below and amplification efficiency was assessed quantitatively by SYBR-based qPCR or qualitatively by gel image band intensity.

#### Emulsion Stabilization Experiments

OIL-PCR test buffer was prepared similarly to the lysis activity assays with ether NEB HF buffer, or Detergent Free Buffer (Thermo Fisher F520L), while omitting bacterial cells or RL Lysozyme. Reactions were emulsified at 25 Hz for 30 seconds on a Retch Mixer Mill MM 400 with adapters 11990 and 11993 (Mobio/Qiagen). Emulsion tubes were photographed before and after thermocycling and assayed by eye for coalescence. After confirming a stable emulsion, qPCR and lysis time series experiments were repeated to confirm activity of Phusion DNA polymerase and RL Lysozyme in the DF buffer.

### OIL-PCR in Tube Based Format for Master Mix Optimization

#### Fusion PCR Reaction Setup

All steps were performed on ice or in a 4°C centrifuge until after emulsification. 50 μl PCR reactions were prepared in a 1.5 ml microcentrifuge tube with varying experimental conditions. 2 μl of bacterial cells standardized to 10^4^ cells/μl were added to 48 μl of master mix and vortexed to evenly disperse cells before adding 300 μl of cold Droplet Generation Oil for Probes (BioRad 1863005). Emulsions were formed immediately after by shaking tubes at 25 Hz for 30 seconds on a Retch Mixer Mill MM 400 with adapters 11990 and 11993 (Mobio/Qiagen). Next, the emulsion mix was divided into four 70 μl aliquots in a PCR strip-tube and thermocycled as follows: 37 °C for 5 minutes, 95 °C for 10 minutes, then 38 cycles of 95 °C for 5 seconds, 54 °C for 30 seconds, and 72 °C for 30 seconds, followed by final extension 72 °C for 2 minutes. After PCR amplification, the aliquots were briefly vortexed and pooled into a clean 1.5 ml microcentrifuge tube. To break the emulsion, 50 μl of TE and 70 μl of Perfluorooctanol (Krackeler Scientific 45-370533-25G) were added and the mixture was vortexed vigorously for 30 seconds. Tubes were centrifuged at 5000 G for 1 minute and the upper aqueous phase was transferred to a new PCR strip tube and purified using AMPure XP beads as described below.

#### Manual AMPure XP Bead cleanup

AMPure XP beads (Beckman A63880) were added at a ratio of 0.8 μl beads per 1 μl of recovered DNA, vortexed, and incubated for 5 minutes for DNA binding. PCR strip tubes were transferred to a 96-Well magnet (Eppendorf Magnum FLX) to pull down beads for 5 minutes. Supernatant was removed with a multichannel pipette and the pellet was washed twice with 100 μl of 70% EtOH before drying for 10 minutes at room temperature. The bead pellet was suspended in 20 – 50 μl of TE and incubated for 5 minutes to elute DNA, before returning to the magnet and transferring supernatant to fresh PCR strip tubes. Eluted DNA was either run directly on a gel for qualitative analysis of amplification, or used as template in qPCR assays.

### Probe-Based qPCR with TaqMan Probes for Cell Input Optimization and Multiplexing

#### Standardization of Bacterial Test Strains

For all experiments, bacterial type strains *Escherichia coli* MG1655^19^, *Vibrio cholerae* N16961^31^, and *Bacillus subtilis* 168^32^ were inoculated from frozen glycerol stocks into 5 ml LB and grown at 37 °C overnight. Cultures were diluted 1:100 in 5ml fresh LB the next day and grown to OD_600_ 0.4-0.8. CFU/μl at OD_600_ was quantified by serial dilution of cells in LB, plating, and colony counting. Count results were used to standardize cell cultures to a stock concentration of 10^6^ CFU/μl to be diluted and used as input for OIL-PCR.

#### Optimizing Cell Input Concentration

Cultures of WT *V*. cholerae N16961^31^ and E. *coli* MG1655^19^ carrying plasmid pBAD33^20^ with *cmR* were standardized to 10^4^, 10^5^, and 10^6^ CFU/μl in LB. The two strains were mixed 1:1 at each of the three concentrations, and 2 μl of cells was used as template in 50 μl tube-based OIL-PCR reactions with fusion primers targeting the *cmR* gene. Reactions with each strain emulsified individually were run as controls. Droplet coalescence was assayed by mixing individual control reactions after emulsification, thereby ensuring the two strains were not encapsulated together. Reactions were thermocycled and recovered DNA was used as template in the probe-based qPCR to quantify specific vs non-specific fusion products.

#### Probe-Based qPCR Assay

Probes were designed to target unique regions of *E. coli* and *V. cholerae* 16S ribosomal rRNA gene. Both probes hybridized to the antisense strand and can only be cleaved when the polymerase extended from the nested *cmR* primer, across the fusion junction, and into the 16S gene, thus distinguishing actual fused PCR products from stray fragments of 16S DNA. Probes were verified to only target their specified strain, with the *V. cholerae* probe having a VIC/NFQ MGB reporter probe and *E. coli* a FAM/NFG MGB probe. 20 μl qPCR reactions were prepared in duplicate as follows: 1x Luna Universal Probe qPCR Master Mix (NEB M3004L), 300 nM of forward and reverse primer, 3.2 μM of forward and reverse blocking primers, 200 nM of *E. coli* and *V. cholerae* TaqMan probes, and 2-5 μl of recovered OIL-PCR amplicons. Reactions were amplified under the following conditions: 95 °C for 1 minutes, then 50 cycles of 95 °C for 20 seconds, 55 °C for 20 seconds, and 60 °C for 20 seconds. For analysis, Ct values were subtracted from the total number of cycles for easier interpretation.

#### Primer Multiplexing Validation

Four strains of bacteria were used to test primer multiplexing: *V. cholerae* N16961^31^ carrying *ampR* on RP4 plasmid^33^, WT *V. cholerae* N16961, *E. coli* 0006 (CDC & FDA Antibiotic Resistance Isolate Bank) carrying *bla*_CTX-M-15_, and WT *E. coli* MG1655^19^, all mixed at a ratio of 1:49:10:40 with a final concentration of 10^4^ cells/μl. This mix of cells resulted in 10% of the consortium carrying *bla*_CTX-M-15_ and 1% carrying *amr*R to provide a more realistic depiction of the abundances of ARGs in natural stool communities. OIL-PCR was performed in a plate-based format with forward and fusion primers for both *amrR* and *bla*_CTX-M-15_. Each strain was tested individually as controls. Purified fusion products were assayed for correct fusions using the probe-based qPCR assay with nested primers targeting *amrR* or *bla*_CTX-M-15_ in parallel reactions.

### Final OIL-PCR Parameters

#### OIL-PCR

The final, optimized OIL-PCR master mix is as follows: 100 U/μl Phusion Hot Start Flex DNA Polymerase, 1X DF Buffer (Thermo Fisher F520L), 250 μM dNTPs (NEB N0447L), 2 μM universal 16S reverse primer 786R, 1 μM of each target specific forward primer, 0.01 μM of each target specific fusion primer with universal 519F’ tail, 1.5 mM additional MgCl_2_, 5mM Ammonium Sulfate, 5 mM DTT, 4 mg/ml BSA (NEB B9000S), 300 U/μl RL Lysozyme, 400 cells/μl Nycodenz purified cells. 300 μl emulsion oil (BioRad 1863005) was added to 50 μl reactions when performed in individual tubes, or 200 μl of emulsion oil was added to 100 μl OIL-PCR reactions when performed in the 96-well plate format. Tubes were emulsified at 25 Hz for 30 seconds, while plates were sealed with a 50 µm aluminum seal (Axygen PCR-AS-600) and emulsified for 2 rounds of 27.5 Hz for 20 seconds; flipping the plate in between for consistent emulsion across rows.

The lysis and amplification program is: 37 °C for 5 minutes, 95 °C for 10 minutes, then 38 cycles of 95 °C for 5 seconds, 54 °C for 30 seconds, 72 °C for 30 seconds, before final extension of 72 °C for 2 minutes.

#### dsDNase Treatment and Heat Inactivation in OIL-PCR

dsDNase treatment was not used in the initial OIL-PCR optimization or spike-in experiments. Cells were standardized to 10^4^ cells/μl in 100 μl of PBS. 1 μl of stock dsDNase (Thermo Fisher EN0771) was added to the tube and incubated at room temp for 10 minutes before returning to ice. Treated cells were used directly in OIL-PCR. The enzyme was inactivated immediately after emulsification (optional) by incubating 10 minutes in a water bath set exactly to 50 °C with gentle mixing by hand every 2 minutes.

#### Nested PCR

SYBR-based qPCR was performed on purified fusion PCR products to minimize the number of cycles for each reaction with the goal of reducing chimera formation. Amplification was performed in 20 μl reactions using the Luna Universal qPCR Master Mix (NEB M3003L) with 1x PCR master mix, 300 nM forward and reverse primers, and 2-5 μl of purified template. For multiplexed experiments, separate reactions were prepared, with one set of nested primers for each gene assayed. The following thermocycling conditions were used: 95 °C for 2 minutes, 40 cycles of 95 °C for 15 seconds, 55 °C for 15 seconds, 68 °C for 20 seconds, followed by a final extension phase at 68 °C for 1 minutes. Melt curves were measured by heating to 95 °C at 0.15 °C /s. Blocking primers were not included in SYBR-based qPCR reactions because of the strong signal from self-hybridization. Ct values were used to select the cycle number for nested amplification that was equal to the Ct value +/- 2 cycles. Reactions that did not amplify in the qPCR were amplified with the highest number of cycles for that preparation.

Using the qPCR results to select the cycle number, nested PCR reactions were prepared in duplicate 20 μl reactions as follows: 20 U/ml Phusion DNA polymerase, 1x HF Buffer, 2 μM dNTPs, 300 nM target gene specific forward primer and universal reverse primer, 32 μM of each blocking primer, and 2-5 μl of template. Thermocycling was performed with variable number of cycles based on the qPCR as follows: 98 °C for 3 minutes, then variable cycles of 98 °C for 5 seconds, 55 °C for 30 seconds, and 72 °C for 30 seconds, followed by final extension 72 °C for 5 minutes. Duplicate PCR reactions were pooled and purified using automated AMPure XP cleanup.

#### Illumina Indexing PCR and Library Preparation

Custom indexing primers were designed based on Spencer et al^11^. A set of unique, 9 bp barcodes was generated using Barcode Generator V2.8^34^. The primers are compatible with the Illumina Truseq primers and the index can be read with 8 bp instead of 9 to make them compatible with other libraries.

Indexing PCR was performed with 25 μl reactions as follows: 20 U/ml Phusion DNA polymerase, 1x HF Buffer, 2 μM dNTPs, 100 nM of unique forward and reverse indexing primers, and 2 μl of purified nested PCR template. Cycling was performed as follows: 98 °C for 1 minutes, then 20 cycles of 98 °C for 15 seconds, 56 °C for 30 seconds, and 72 °C for 45 seconds, followed by final extension 72 °C for 2 minutes. PCR reactions were purified using automated AMPure XP cleanup.

Indexed PCR libraries were quantified using QUANT-IT pico green dsDNA assay kit (Invitrogen P7589) and measured on the Spectramax M3 plate reader. Wells were pooled based on the measured concentration using the Eppendorf epMotion 5075vtc robot and the final pool quantified using the Qubit Broad Range Assay Kit (Thermo Fisher Q32853). Pools were run on a gel to confirm clean DNA before sequencing with MiSeq 2×250 V2 chemistry.

### Plate-Based OIL-PCR With Robotic Automation

#### Reaction Setup

96 μl of the final OIL-PCR master mix was aliquoted into a 500 μl deep well plate (Eppendorf 00.0 501.101). Nycodenz purified stool cells were diluted in PBS to 10^4^ cells/μl in an 8-well PCR strip for multichannel pipetting. 4 μl of cells was quickly added to the reactions with a 10 μl 8-channel pipette before sealing with an extra-thick foil seal (Axygen PCR-AS-600) and vortexed to mix. The reactions were briefly centrifuged to return liquid to the bottom of the plate, and then placed on an orbital microplate shaker (VWR 12620-926) at 1200 rpm for 30 seconds to further mix the cells while keeping the mix at the bottom of the wells. After mixing, the foil seal was carefully removed and 200 μl of cold emulsion oil was added using a multichannel pipette. The plate was then sealed with a fresh foil seal and shaken at 27.5 Hz for 20 seconds on the Retch shaker MM 400 with plate adapter (#11990). The plate was removed and turned over to shake an additional 20 seconds providing an even emulsion across the plate. After emulsifying, each reaction was aliquoted into 4 wells of a PCR plate (Eppendorf 0030 128.648) using the robot for consistency. The plates were sealed and run on the OIL-PCR fusion program described earlier.

#### DNA Recovery from Emulsion

After amplification, the robot was used to purify the OIL-PCR Products. In short, replicate reactions were pooled into a fresh 500 μl deep well plate, and 60 μl of TE and 70 μl of Perfluorooctanol (Krackeler Scientific 45-370533-25G) were added to each well. The plate was sealed and shaken on the Retch at 30 Hz for 40 seconds to thoroughly disrupt the emulsion. The plate was then centrifuged in a swing bucket rotor at 5000 Gs for 1 minute to separate the phases and returned to the robot. 80 μl of the upper phase was aspirated from a defined height into a fresh 500 μl deep well plate for automated Ampure XP bead purification.

#### Automated AMPure XP Bead Purification

85 μl of AMPure XP beads (Beckman A63880) was added to the deep well plate containing the recovered OIL-PCR fusion products. The reactions were mixed at 1200 rpm for 1 minute and incubated for 2 minutes for DNA binding, before transferring to the magnet (Eppendorf Magnum FLX) for 3 minutes. After pulldown, the supernatant was discarded and the wells were washed twice with 200 μl of 70% EtOH. After discarding the second wash, the plate was removed from the magnet and dried at room temp for 10 minutes before adding 50 μl of TE buffer. The plate was shaken at 1200 rpm for 1 minute and incubated 2 minutes to elute the DNA. Finally, the plate was returned to the magnet and for 2 minutes and 48 μl of purified DNA was transferred to a fresh 96-well PCR plate (Eppendorf 0030 128.648).

### OIL-PCR on Natural Stool Communities

#### Nycodenz Purification of Stool Cells

All steps were performed on ice and in a 4c refrigerated centrifuge unless otherwise noted. Stool samples were collected in PBS + 20% glycerol + 0.1% L-cysteine and frozen at -80c until processed. Frozen samples were thawed completely and thoroughly homogenized via vortexing. Samples were diluted at least 1:1 in cold PBS to reduce the sample viscosity and glycerol concentration as viscous samples did not to separate well with the Nycodenz. Samples were vortexed at maximum speed for 5 minutes to release cells from stool particles. 300μl of cold 80% Nycodenz (VWR 100356-726) was aliquoted to the bottom of 2 ml microcentrifuge tubes and 1.6 ml of stool slurry was overlaid on top without mixing the two phases. Tubes were centrifuged at 10,000 G for 40 minutes in a swing bucket rotor to separate cells. After centrifugation, the upper phase was removed with a pipetted and 500μl of cold PBS was used to wash the bacterial cell pellet from the insoluble stool fraction. The suspended cells were removed and the pellet was washed a second time with 500 μl of PBS. Cells were centrifuged at 50g for 1 minute to pellet any large particles that carried over from the Nycodenz purification and the upper phase was passed through a 40 μm nylon mesh screen (Falcon 352235) to remove any residual stool debris or large cell clumps. Samples of each preparation were diluted 1:1 PBS + 20% glycerol for whole cell storage. Lastly, purified cells were diluted and imaged at 100x magnification within a 20 μm counting chamber (VWR 15170-048). Images were analyzed using FIJI/ImageJ 1.52p (Java 1.8.0_172) to manually count cells and calculate cell concentration in glycerol stocks.

#### Spike-In Experiment

This experiment was performed using the individual tube-based format of OIL-PCR. Nycodenz purified stool and *E. coli* carrying pBAD33^20^ with *cmR* was standardized to 10^4^ cells/μl. *E. coli* cells were mixed with the stool samples at a ratio of 1:10, 1:100, and 1:1000, and the mixed cultures were added to OIL-PCR containing the *cmR* primer set. Reactions were emulsified, lysed, thermocycled, and fusion products were purified manually. Nested PCR was performed with the nested *cmR* primer before indexing, pooling, and sequencing.

#### Lysozyme, dsDNase, Heat Experiment

Nycodenz-purified human and chicken stool cells were standardized to 10^5^ cells/μl and incubated with or without dsDNase at room temp for 10 minutes. OIL-PCR master mix was prepared with and without Lysozyme using universal 16S rRNA primers i519F and i786R. Cells were added to the OIL-PCR reaction and emulsified. Emulsions were either incubated at 50 °C or room temperature for 10 minutes before aliquoting to PCR plates and running the OIL-PCR fusion program. Amplicons were purified, indexed, and submitted for Illumina sequencing as described above.

### OIL-PCR for Detection of *bla* genes in Neutropenic Patients

#### Sample collection and Metagenomic Assemblies

Samples were collected, sequenced, and metagenomic assemblies were prepared as described in Kent et al.^2^ Briefly, serial stool samples were collected from consenting individuals receiving a hematopoietic stem cell transplant at NewYork-Presbyterian Hospital/Weill Cornell Medical Center in accordance with IRB protocols for Weill Cornell Medical College (#1504016114) and Cornell University (#1609006586). Samples were either frozen “as is” (for metagenomic sequencing) or homogenized in phosphate-buffered saline (PBS) + 20% glycerol before freezing (for OIL-PCR). DNA was isolated from samples destined for metagenomic sequencing using the PowerSoil DNA Isolation Kit (Qiagen) with additional proteinase K treatment and freeze/thaw cycles recommended by the manufacturer for difficult-to-lyse cells. Extractions were further purified using 1.8 volumes of Agencourt AMPure XP bead solution (Beckman Coulter). DNA was diluted to 0.2 ng/μL in nuclease-free water and processed for sequencing using the Nextera XT DNA Library Prep Kit (Illumina).

#### Design and Validation of OIL-PCR fusion primers

ARG variants for the three *bla* genes were downloaded from the CARD database^35^ and aligned in Snapgene using default MUSCLE parameters. Conserved regions were identified manually and degenerate primers were designed to capture as many variants of the genes as possible. Primers were selected for GC content between 40 – 60% and an annealing temperature of 58 °C based on the Snapgene calculation. Degenerate bases were limited to 3 per primer and no less than 5 bp from the 5’ end.

Strains acquired through the CDC & FDA Antibiotic Resistance Isolate Bank carrying multiple variants of each gene (Supplementary Table T1)) were used as template for testing *bla* primers. At least 3 sets of primers were designed and tested in every possible combination using the OIL-PCR master mix without emulsion to find a set of three primers that provided clean fusion amplification. Lastly, working primer sets were tested in an emulsion on whole cells to confirm amplification in OIL-PCR.

Fusion Primers targeting Tn3 transposon genes were designed using scaffolds from the metagenomic assemblies and tested on *Klebsiella* isolate DNA from patient B335.

#### OIL-PCR on Neutropenic Patients

All OIL-PCR reactions were performed with the plate-based protocol including dsDNase treatment and heat inactivation. Whole bacterial cells were purified with Nycodenz, quantified and standardized to 10^4^ cells/μl in PBS before treating with dsDNase. For multiplexed experiments, reactions were prepared in quadruplicate with three sets of primers targeting the three *bla* genes in each reaction. The singleplex reactions were prepared in triplicate with only one primer set per reaction. In all cases, the reactions followed the standard plate-based protocol with automation, including heat inactivation of the dsDNase after emulsification. Nested PCR, indexing, and library preparation was performed as described above.

#### Romboutsia Specific OIL-PCR

CRIB primers^25^ were modified to form a fusion product with all three *bla* genes, however, only the *bla*_TEM_ primer set amplified when tested. Using only the *bla*_TEM_ primer set, OIL-PCR was performed with dsDNase treatment, in triplicate, using the plate-based format with automation. Nested PCR, indexing, and library preparation was performed as described above.

### Computational Methods

#### Processing 16S rRNA Sequencing

Raw reads were merged using usearch^36^ (V 11.0.667) -fastq_mergepairs (maxdiffs: 20, pctid: 85, minmergelen: 283, maxmergelen: 293) before trimming primers and quality filtering with usearch - fastq_filter (maxee: 1.0). Unique reads were filtered using usearch -fastx_uniques and OTUs were clustered based on 97% identity with usearch -cluster_otus. OTU tables were generated with usearch - otutab and taxonomy was assigned with RDP classifier implemented in MOTHUR classify.sequs (1.38.1) against silva v132. Rarefaction curves were generated using QIIME1^37^ (v1.9) multiple_rarefaction.py (-m 10, -x 100000, -s 100, -n 5, -k).

#### Processing OIL-PCR Sequencing

Raw reads were merged using usearch^36^ (V 11.0.667) -fastq_mergepairs (maxdiffs: 10% of expected overlap, pctid: 85, minmergelen: expected length-15, maxmergelen: expected length +15) before trimming primers and quality filtering with usearch -fastq_filter (maxee: 1.0). Unique reads were filtered using usearch -fastx_uniques. Reads were split at the fusion junction into 16S and target reads using cutadapt V2.1^38^ because of its tolerance for PCR errors which are often introduced in the fusion junction of the OIL-PCR amplicons. The 16S reads were clustered based on 97% identity with usearch - cluster_otus, OTU tables were generated with usearch -otutab and taxonomy was assigned with RDP classify implemented in mothur^39^ classify.sequs (1.38.1) against SILVA^40^ v132. Target reads were identified by blasting against a custom database of expected sequences with blastn^41^ (v2.9.0). 16S taxonomy and target read identity were then reassociated using a custom python script to parse the files. Detections were defined by taxa – target associations that make up 0.5% of the total reads across replicates.

#### Strain Level Analysis of *Romboutsia* in Metagenomes

Metagenomic reads from each time point were aligned to the R. timonensis reference genome (Refseq accession code: GCF_900106845.1) using BWA mem (v0.7.17, -a)^42^. Reads aligning to the reference genome were then assembled using SPAdes (v3.14.)^43^. To determine the presence and identity of strains from each time point, AMPHORA^44^ (v2, marker identification step only) was used to identify the sequences of 30 marker genes within each assembled *R. timonensis* genome. The marker genes identified by AMPHORA were then mapped (Diamond, v2.0.4)^45^ to the BLAST^46^ nr database for taxonomic annotation (BLAST nr database downloaded 2018). DNA sequences of the marker genes that mapped to *R. timonensis* were retained for further analysis. Genes from time point 2 and time point 2 were aligned to one another (BLAST blastn, v2.9.0)^41^, and then sequences from time point 1 were aligned against sequences of the same gene from time point 2, once the sequences at time 2 and time 3 were determined to be the same. To determine how abundant each marker gene, and all of its variants, are at each time point, metagenome reads from each time point were mapped to its own set of marker gene sequences (BWA mem, v0.7.17, -a)^42^. Read counts were normalized for the length of each gene and the total number of reads sequenced per sample (RPKM)^47^.

## Supporting information

Table1_Strains_adn_Plasmids

Table2_Primers

Table3_OIL_Primer_Sets

## Acknowledgments

We would like to thank the following individuals and organizations for their generosity in providing us with strains and plasmids: Tobias Dörr (STRAINS), Barth Smets (RP4), John Helmann (*B. subtilis*) and the CDC & FDA Antibiotic Resistance (AR) Isolate Bank. We would like to thank Sarah Spencer for technical advice. This study was funded by the Centers for Disease Control (OADS BAA 2016-N-17812) and by the National Sciences Foundation (Awards #1661338 and #1650122). F.N. is a SUNY Diversity Fellow. M.J.S. is funded by the NIAID (K23 AI114994). I.L.B. is funded by the NIH (1DP2HL141007-01) and is a Sloan Foundation Research Fellow, a Packard Fellowship in Science and Engineering, and a Pew Foundation Biomedical Scholar.

